# Dockground scoring benchmarks for protein docking

**DOI:** 10.1101/2021.09.02.458795

**Authors:** Ian Kotthoff, Petras J. Kundrotas, Ilya A. Vakser

## Abstract

Protein docking protocols typically involve global docking scan, followed by re-ranking of the scan predictions by more accurate scoring functions that are either computationally too expensive or algorithmically impossible to include in the global scan. Development and validation of scoring methodologies are often performed on scoring benchmark sets (docking decoys) which offer concise and nonredundant representation of the global docking scan output for a large and diverse set of protein-protein complexes. Two such protein-protein scoring benchmarks were built for the Dockground resource, which contains various datasets for the development and testing of protein docking methodologies. One set was generated based on the Dockground unbound docking benchmark 4, and the other based on protein models from the Dockground model-model benchmark 2. The docking decoys were designed to reflect the reality of the real-case docking applications (e.g., correct docking predictions defined as near-native rather than native structures), and to minimize applicability of approaches not directly related to the development of scoring functions (reducing clustering of predictions in the binding funnel and disparity in structural quality of the near-native and non-native matches). The sets were further characterized by the source organism and the function of the protein-protein complexes. The sets, freely available to the research community on the Dockground webpage, present a unique, user-friendly resource for the developing and testing of protein-protein scoring approaches.

## Introduction

Proteins most often function by interacting with other proteins. Structural characterization of these interactions is important for our ability to understand and modulate them. Experimentally determined structures of protein-protein complexes constitute only a small fraction of the known interactome.^1^ Thus, computational techniques to model three-dimensional structures of protein-protein complexes (protein-protein docking) are required to fill the gap. In recent years, such techniques have been rapidly developing.^2^ They can be roughly divided into free docking and template-based docking approaches. The free docking is performed without *a priori* knowledge of an experimentally determined structure of similar/homologous complexes, while the template-based docking explicitly utilizes such knowledge.^3^ At the initial global search (scan) stage, both free and template-based docking produce putative docking models. The correct (near-native) model can be hidden among them with a relatively low rank. Thus, docking approaches routinely involve a scoring stage, at which protein-protein complexes output from the global scan are re-scored (re-ranked) by more accurate functions that are either computationally too expensive or algorithmically impossible to include in the global scan.

A number of scoring functions have been developed in order to distinguish near-native from incorrect docking predictions.^4-6^ These functions are often tested on non-redundant curated sets of experimentally determined structures of protein-protein complexes (docking benchmarks). Such sets consist of the native or near-native structures of protein-protein complexes composed from the unbound forms of the constituent proteins.^7,8^ Thus, generation of putative docking models is left to the researcher, which may lead to a bias in comparing performance of different scoring functions. To mitigate this problem by providing a standard and convenient testing ground for the community of scoring functions developers, one needs pre-generated sets of near-native and incorrect docking models (docking decoys) for each protein-protein complex in the docking benchmark. Several such sets (scoring benchmarks) already exist. The Dockground unbound decoy set 1 consists of 61 protein-protein complexes, each with 100 docking poses of which at least one is a near-native and the others are decoys.^9^ Sternberg and co-workers built a decoy set for training FTDock program.^10^ CAPRI (Critical Assessment of Predicted Interactions)^6^ scoring benchmark consolidates predicted complexes from the CAPRI community-wide scoring experiment.^11^ Weng lab maintains extensive sets of decoys, based on their protein docking benchmarks,^7,12-15^ generated by ZDOCK^16^ and ZRANK^17^ programs. RosettaDock was used to generate docking decoys based on flexible docking.^18^ Some of these decoy sets do not have a near-native match.

Existing decoy sets focus on experimentally determined structures. To address limited structural accuracy of protein models, especially in high-throughput (e.g. genome-wide) modeling, we developed docking benchmarks composed of models of the individual proteins. For each protein we generated several models of different accuracy assessed either by the model’s RMSD from the native structures (Dockground model-model docking benchmarks 1^19^ and 2^20^) or by the rank from structure prediction software (model-model docking benchmarks Q1 and Q2^21^). In this paper, we present two new large scoring benchmark sets (docking decoys). A decoy set of experimentally determined unbound structures from the docking benchmark 4^8^ and a decoy set of protein models from the model-model docking benchmark 2. The decoy sets are publicly available in the Dockground resource at http://dockground.compbio.ku.edu.

## Results and Discussion

### Design principles

Docking decoy sets (scoring benchmarks) should reflect the reality of the real case scenario docking applications. At the same time, they should be hardened agains simple ways to “defeat” them by trivial approaches not directly related for the development of scoring functions. Thus, we applied following general principles to the design of adequate scoring benchmarks.

#### Near-native vs. native

A decoy set should contain a few *correct* docking poses and many *incorrect* ones. While it is tempting to define the correct pose as the native (i.e. esperimentally determined) structure, in practice, actual native poses are almost never predicted by the docking procedures. Thus their inclusion into the decoy set is unrealistic, and the correct docking should be defined as a *near-native* pose. A docking pose can be determined as near-native according to establised in the community criteria (e.g. CAPRI criteria^22^). Also, importantly, since near-native prediction is not positioned at the very bottom of the intermolecular energy funnel,^23^ it is more difficult to distinguish from the non-native/incorrect ones, which again makes the set more adequate to the real-case docking scenario.

#### Similar scores

In typical global docking scan output, the correct (near-native) prediction would be placed down the ranked list of predictions, below a number of the incorrect predictions. The number of the incorrect predictions can be large, especially in the free docking (e.g. tens of thousands, or more). This problem is the entire reason for the development of the scoring functions to improve the ranking of the predictions. A simple and thus tempting way to construct a decoy set is to select the top ranked incorrect matches (e.g., ranks 1-99), and combine them with the highest ranked near-native match (e.g., rank ∼100,000), which was a common approach in earlier docking decoy sets. A trivial way to “defeat” such set is to ask the procedure to look for the *worst* (in terms of energy, interface area, etc.) rather than the *best* match. Not surprisingly, such approach almost always correctly identifies the near-native solution in such decoy sets. To avoid even an implicit influence of such line of inquiry, the incorrect docking matches need to have global scan scores similar to the ones of the near-native poses.

#### Clustering

The intermolecular energy funnel can be detected by clustering of the low-energy docking matches.^24^ Such clustering is a strong indicator of the correct binding mode. However, to develop scoring functions, which are not based on this obvious and simple criterion (and thus can complement it), the scoring benchmarks need to avoid clustering around near-native matches, spatially distributing the non-native poses as uniformly as possible.

### Unbound docking decoys

One of the problems in protein-protein docking is the ability of a method to deal with conformational changes upon protein binding. To develop and test procedures capable of successfully addressing this problem, one needs docking decoys based on the unbound structures of the interacting proteins. We generated such set of docking decoys for each of the 396 unbound-unbound complexes from the Dockground unbound docking benchmark 4. First, 300,000 docking solutions for each complex were produced by GRAMM^25,26^ in rigid-body FFT-based docking mode, with the default grid step 3.5 Å and 10° angular interval. To exclude interference from any post-processing scoring and/or structural refinement, docking poses were unscored, unrefined, and ranked only by the GRAMM’s global scan stage shape complementarity. Because of that, most near-native docking poses were outside top 100,000 predictions (Figure 1). The near-native complexes were defined as acceptable or better according to the CAPRI criteria^22^ calculated with respect to the reference complex of the unbound proteins aligned to the bound structures of the native complex.

**Figure 1.**
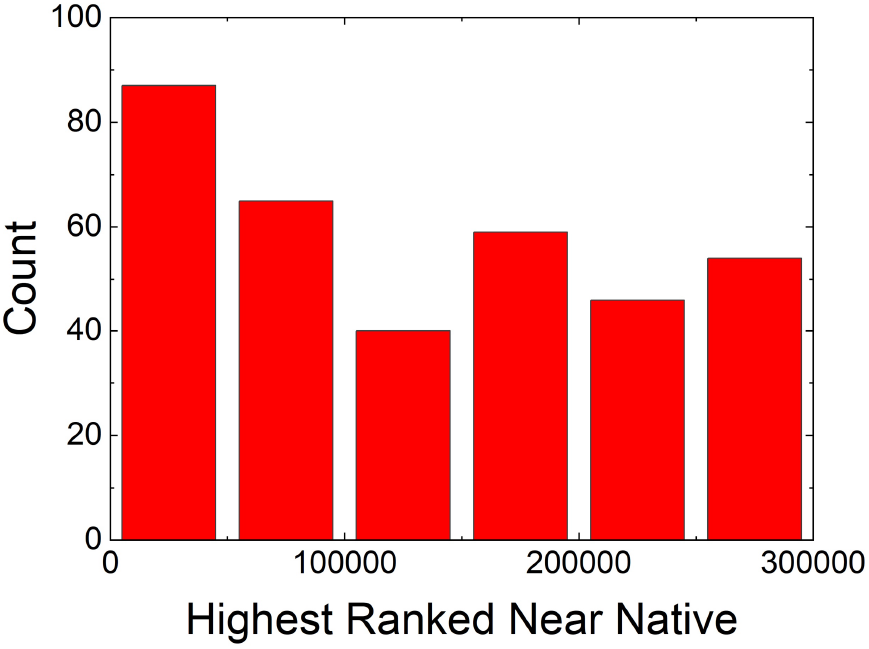
Distribution of unranked near-native poses in the unbound decoy set.

The set contains one near-native solution and 99 docking poses, which are incorrect by the CAPRI classification, selected from 300,000 docking predictions per complex. For each complex, we identified a near-native docking pose ranked highest by the global scan shape complementarity. If such near-native docking pose was absent in the 300,000 docking predictions, the complex was excluded from the set (in our docking experience, complexes without near-native solution in 300,000 predictions are not suitable for rigid-body docking). In order to have similar interface areas (assessed by the global scan shape complementarity) for all docking poses (see the above *Design Principles*), incorrect docking poses were appended to the decoy set initially from the sub-list containing incorrect docking poses within ±50 ranking positions from the near-native match. To reduce spatial clustering (according to the above *Design Principles*), a pose from that sub-list was added to the decoy set only if the angles between the vector connecting geometric centers of the receptor and the ligand and such vectors for the previosuly selected ligand poses were >5°. If this sub-list was exhausted and still less than 99 incorrect docking matches added to the decoy set, the sub-list was expanded by including another 50 lower and 50 higher-ranked matches with the algorithm making another pass through the data. In addition, the minimum angle between poses was reduced by half with each iteration. The protocol iterated until the decoy set was full (containing one near-native and 99 incorrect docking poses).

This decoy set is available for download from the Dockground resource http://dockground.compbio.ku.edu via the decoys page as either the entire set or as individual complexes (Figure 2). The page provides additional information on each decoy set: rank by GRAMM gloabl scan of the highest ranked near-native pose; C^α^ RMSD of the docked ligand; interface C^α^ RMSD; PDB code and chains ID of the bound and the unbound proteins; and bound/unbound C^α^ RMSD.

**Figure 2.**
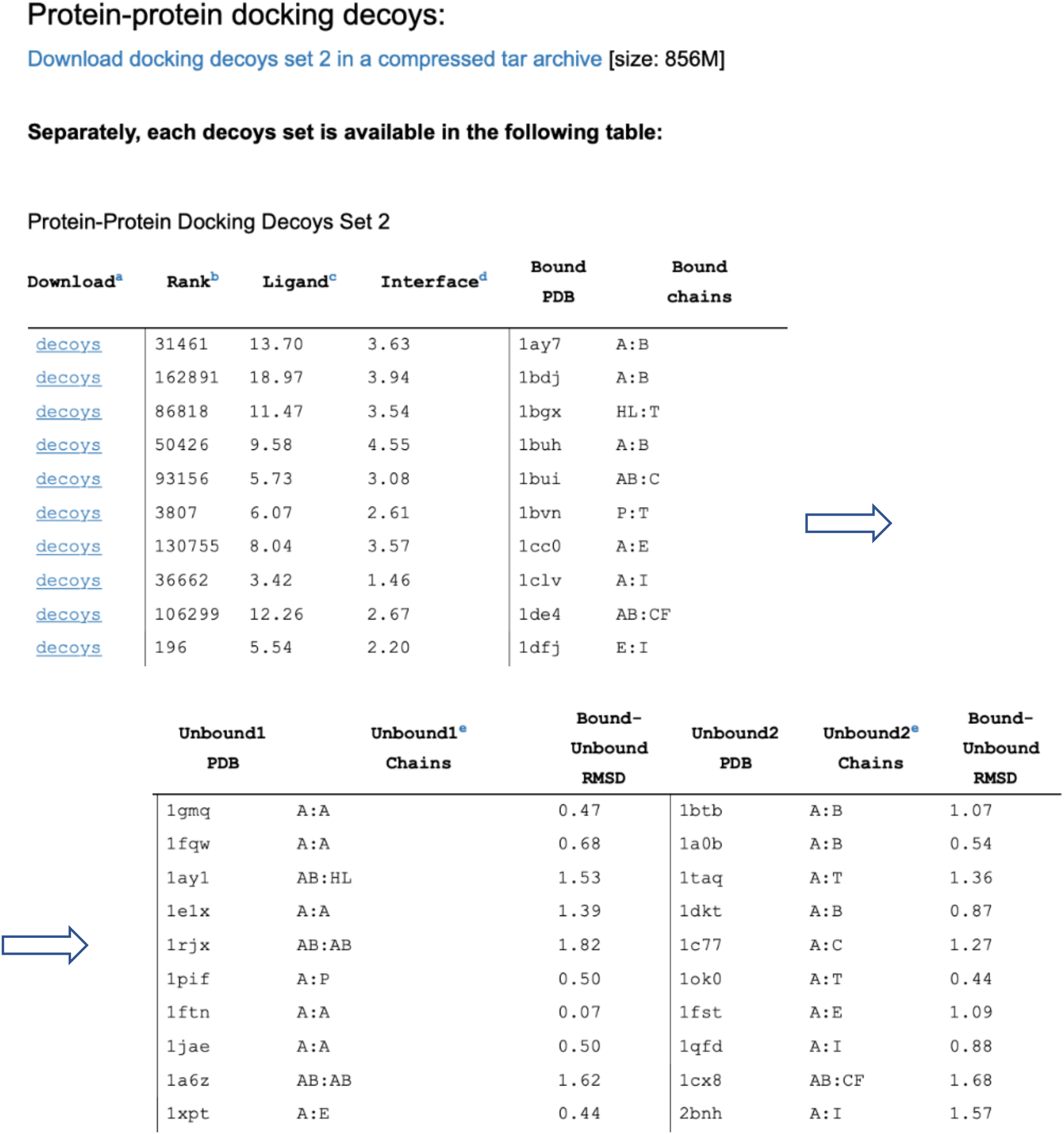
Web interface for the unbound decoy set. Users can download all decoys or a specific decoy set.

### Model docking decoys

The model-model docking decoy set was built from the Dockground model-model docking benchmark set 2^20^ which contains models of individual proteins from 165 protein-protein complexes at pre-determined structure accuracy levels from the experimentally determined structure. The accuracy was defined as C^α^ RMSD between protein in the experimentally determined structure of the complex and the model of that protein (optimally aligned on the native protein). The accuracy levels in the model benchmark set 2, and thus in the docking decoys set, are 1, 2, 3, 4, 5 and 6 Å. The complexes in the model benchmark set 2 have: (*i*) redundancy removed by the sequence identity 30% threshold between pairs of chains, (*ii*) buried solvent accessible surface area >250 Å^2^ per chain, and (*iii*) at least 10 interface residues in each chain.^20^

Docking decoys were generated for each pair of the protein models. For simplicity, we considered only protein pairs with the same accuracy level. Similarly to the unbound docking decoys, we considered 300,000 low-resolution docking solutions produced by GRAMM. The pool consisted of such large amount of putative docking poses because for most protein-protein complexes in the set, the top near-native docking match in the global docking output ranked outside 100,000 solutions (Figure 3). Re-ranking of the docking poses by our AACE18 contact potential^27,28^ significantly improved the ranking of the near-native poses (Figure 3). However, similarly to the unbound decoy set, to avoid interference of the post-processing scoring, we did not use the re-scored improved ranking. Thus, for each level of model accuracy in a complex, the correct pose was designated as the near-native docking match that had the highest ranking according to the global docking output. The 99 incorrect docking poses were selected to have similar shape complementarity scores and spatial distribution avoiding clustering, according to the above approach for the generation of the unbound decoys. The procedure produced spatially well distributed docking poses, as illustrated in Figure 4, at all model accuracy levels. If the near-native docking solution at a given accuracy level was absent in the 300,000 docking poses, the protein-protein complex was excluded from the set.

**Figure 3.**
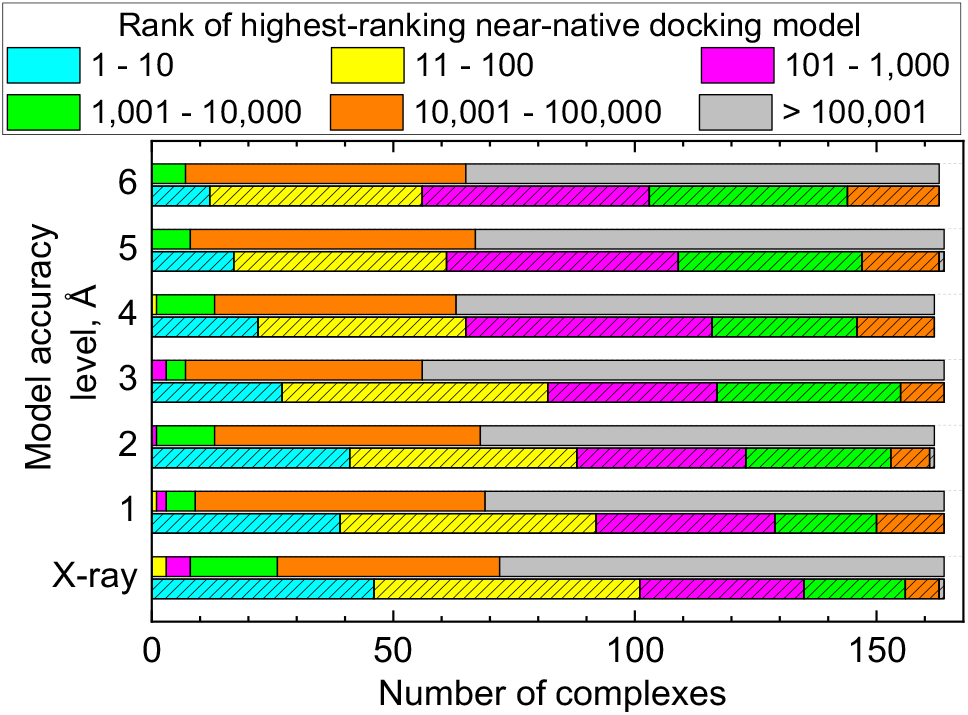
Distribution of docking poses in the model-model docking decoys set. The ranks of the top near-native solution are from the global docking search (open bars) and those scored by AACE18 potentials (hatched bars) for experimentally determined and modeled structures of individual proteins in 165 protein-protein complexes from the Dockground model set 2.

**Figure 4.**
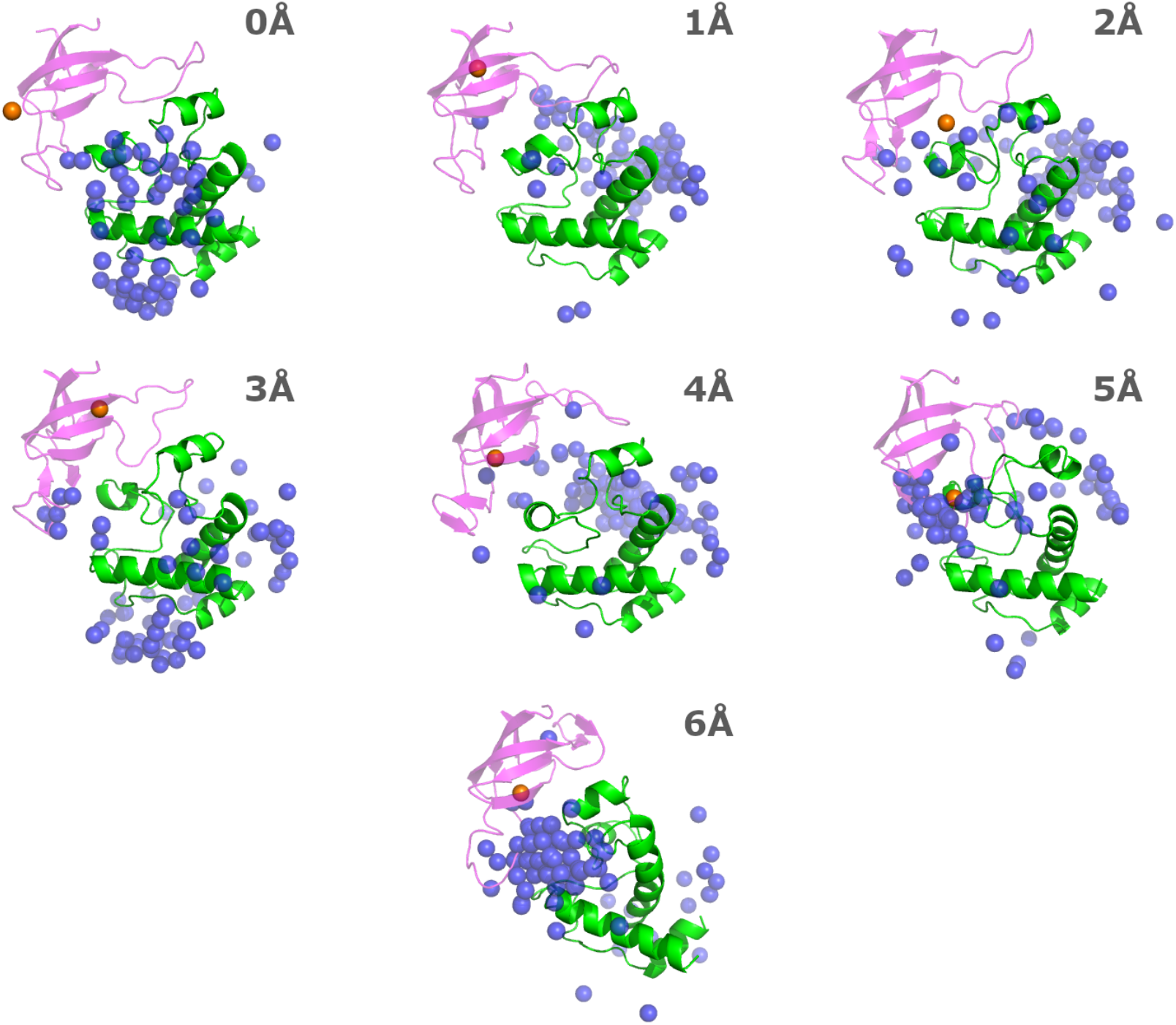
Example of docking decoys set. The proteins are the ferredoxin thioredoxin reductase complex (PDB code 1dj7, chains A and B). The decoys are generated for the experimentally determined structure (0Å) and models of the individual proteins with pre-set accuracy levels 1-6Å. The receptor (chain A) is in green. Spheres are the geometric centers of the ligand (chain B) in incorrect matches (blue) and in the highest-ranking near-native match (orange). The model of the ligand superimposed on the native pose of the ligand in the experimentally determined structure is in magenta.

Decoy sets were successfully generated for 164 of the original 165 experimentally determined protein-protein complexes. For 160 complexes, the decoy sets were generated at all accuracy levels of the individual proteins. For reference, we also generated decoy sets for the native protein-protein complexes. A full list of the successfully generated decoy sets, along with an interactive interface for customizable download, are available on the decoys page of the Dockground resource http://dockground.compbio.ku.edu. An example of the download page is in Figure 5. On this page, one can select to download all decoys, all decoy sets at a specific level of structural accuracy, all levels of accuracy for a protein-protein complex, or a custom selection of the sets.

**Figure 5.**
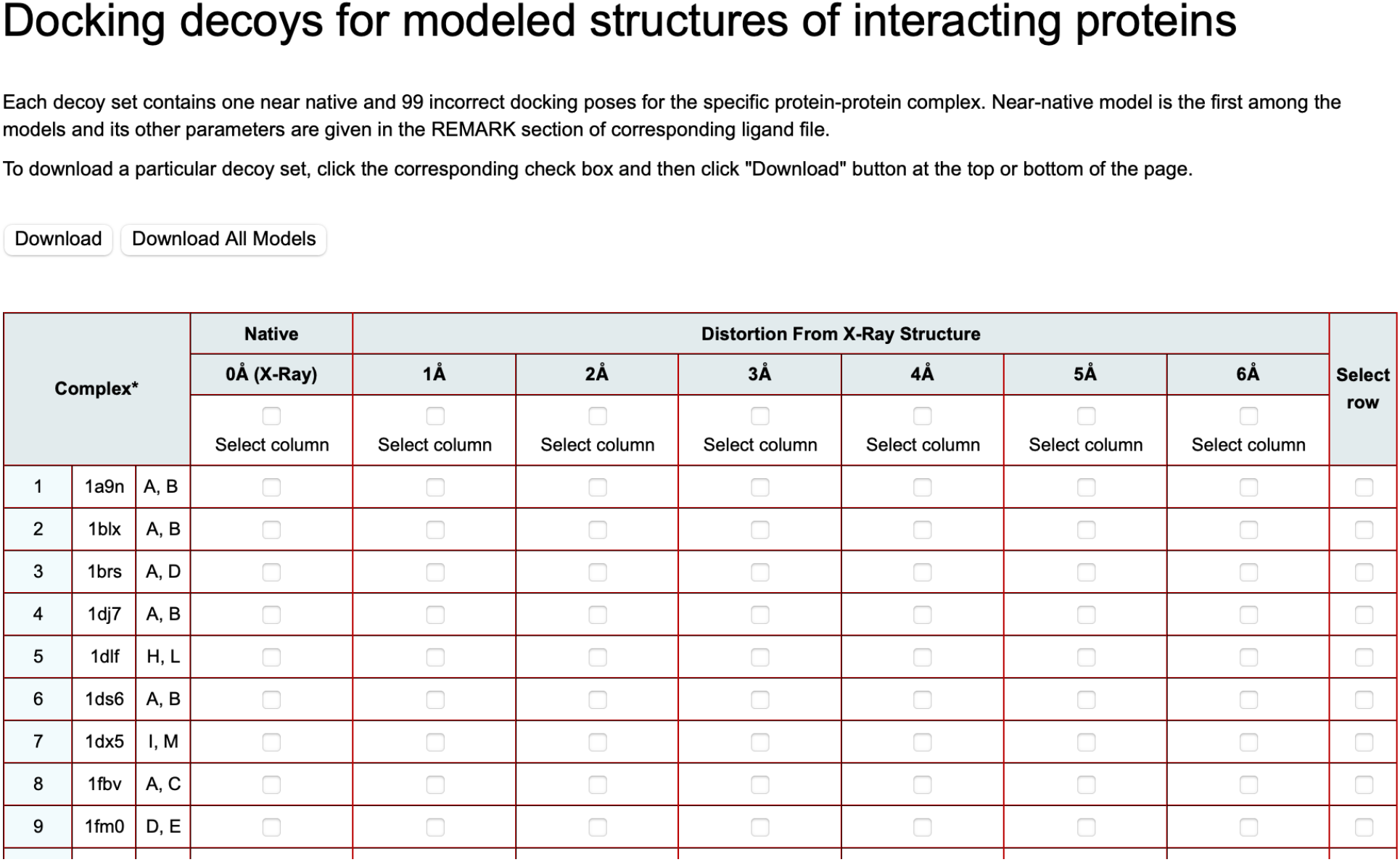
Web interface for model-model docking decoys. Users can select to download all decoys, all decoys at a specific level of accuracy, decoys at all levels of accuracy for a specific complex, any specific decoy set, or a custom selection.

#### Source organisms

In the unbound docking decoy sets, the top three source organisms are: human (103 complexes or 26% of the set have at least one protein from human), *Escherichi*a *coli* (25 complexes or 6%) and mouse (16 complexes or 4%). In the model decoy set, the top three organisms are the same as in the unbound set: at least one protein in 70 complexes (out of 164, or 43%) is from human, with the next two largest source species *Escherichi*a *coli* (at least one protein in 14 complexes or 9%) and mouse (13 complexes, or 8%). In the entire PDB, 25% of binary complexes have, at least, one protein from human, also followed by *Escherichi*a *coli* (5%), and mouse (3%). Due to computational constraints, in the entire PDB we considered only binary complexes, i.e. structures with two polypeptide chains in both asymmetric and biological units. Each of these organisms are commonly used as a test system for their respective phylogenic group and as such is expected to be well represented in PDB and its subsets.

Within broader categories of the source organisms, most complexes in both unbound and model docking decoys sets have at least one protein from higher eucaryotes (162 and 104 complexes from unbound and model sets, respectively), followed by bacteria (62 and 29 complexes), lower eucaryotes (13 and 8 complexes), viruses (14 and 8 complexes) and archaea (4 and 6 complexes). This order is similar in the entire PDB with the exception of bacteria, which has greater representation in PDB than every other group (Figure 6).

**Figure 6.**
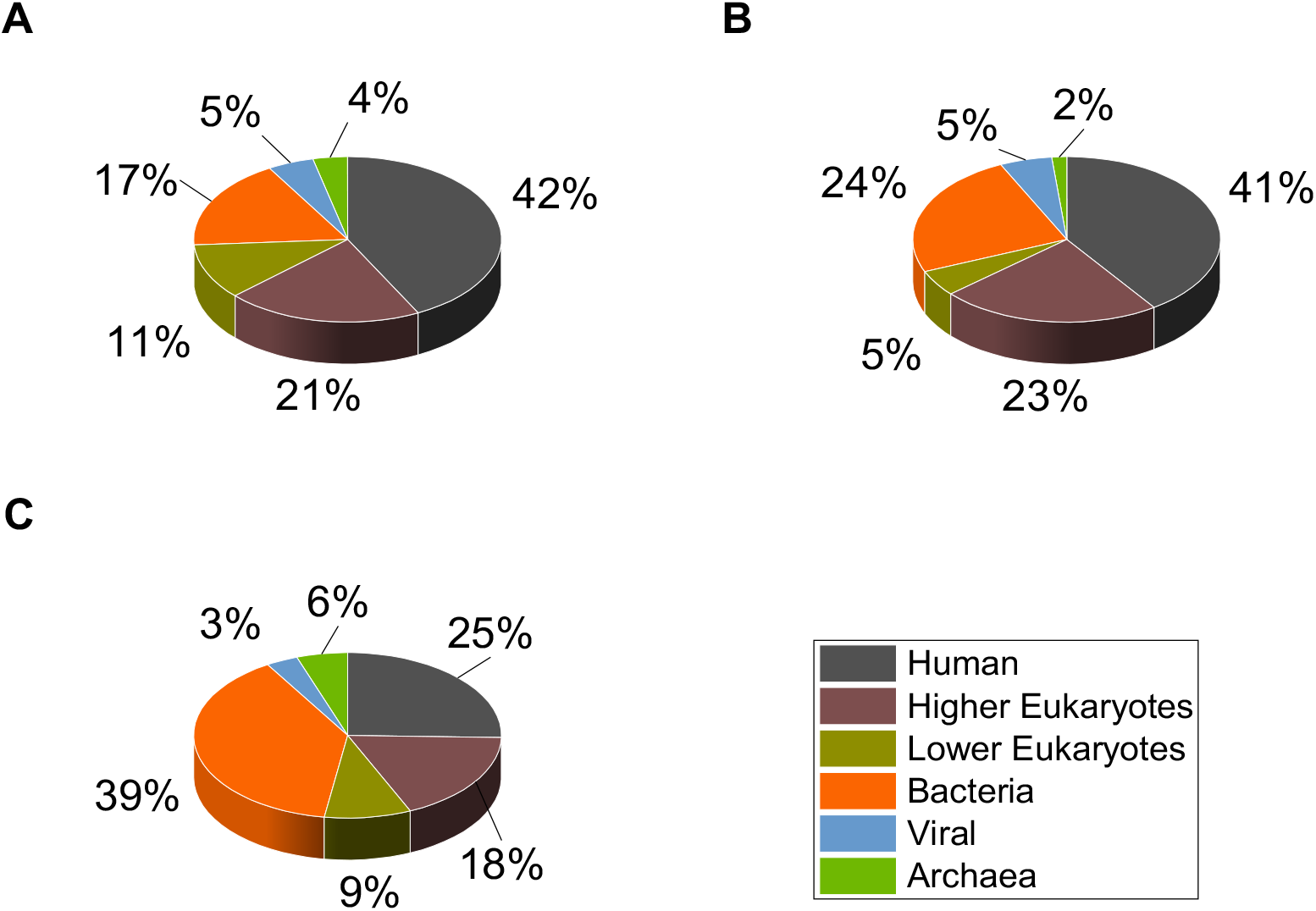
Datasets source organisms. The statistics are for protein-protein complexes in (A) unbound and (B) model docking decoy sets, and (C) for all binary complexes in PDB.

### Functional annotations

The docking decoys sets were further characterized by the function of the protein complexes. This was done using gene ontology (GO) terms, which are subdivided into three domains - molecular function, biological process and cellular component. The trems are organized in a directed acyclic graph with several types of connections between them (we considered only “is-a” type, i.e. “parent-child” relationship). To put complexes in categories large enough for statistical significance, but not characterized by GO terms that are too generic, we utilized GO terms at level two of the molecular function graph. In the unbound docking decoy set, the most common GO term was hydrolase activity (61 protein-protein complexes). It was followed by organic cyclic compound binding and catalytic activity acting on a protein (19 complexes), protein binding (18 complexes), and organic substance metabolic process (15 complexes; Figure 7A). The most common GO terms in the model docking decoy set were hydrolase activity, organic substance metabolism and organic cyclic compound binding - each occurring at least once in 16 protein-protein complexes, followed by catalytic activity acting on a protein (12 complexes), and protein binding (8 complexes; Figure 7B). Although the order is different in the unbound and the model sets, the top five most common functions are the same. GO terms were also determined for the binary complexes from the entire PDB (Figure 7C). Two of the top five most common GO terms from both decoy sets, hydrolase activity and catalytic activity acting on a protein, are also present in the top five categories in entire PDB binary set. The other three are all within the top 20 most common GO terms of the binary PDB set.

**Figure 7.**
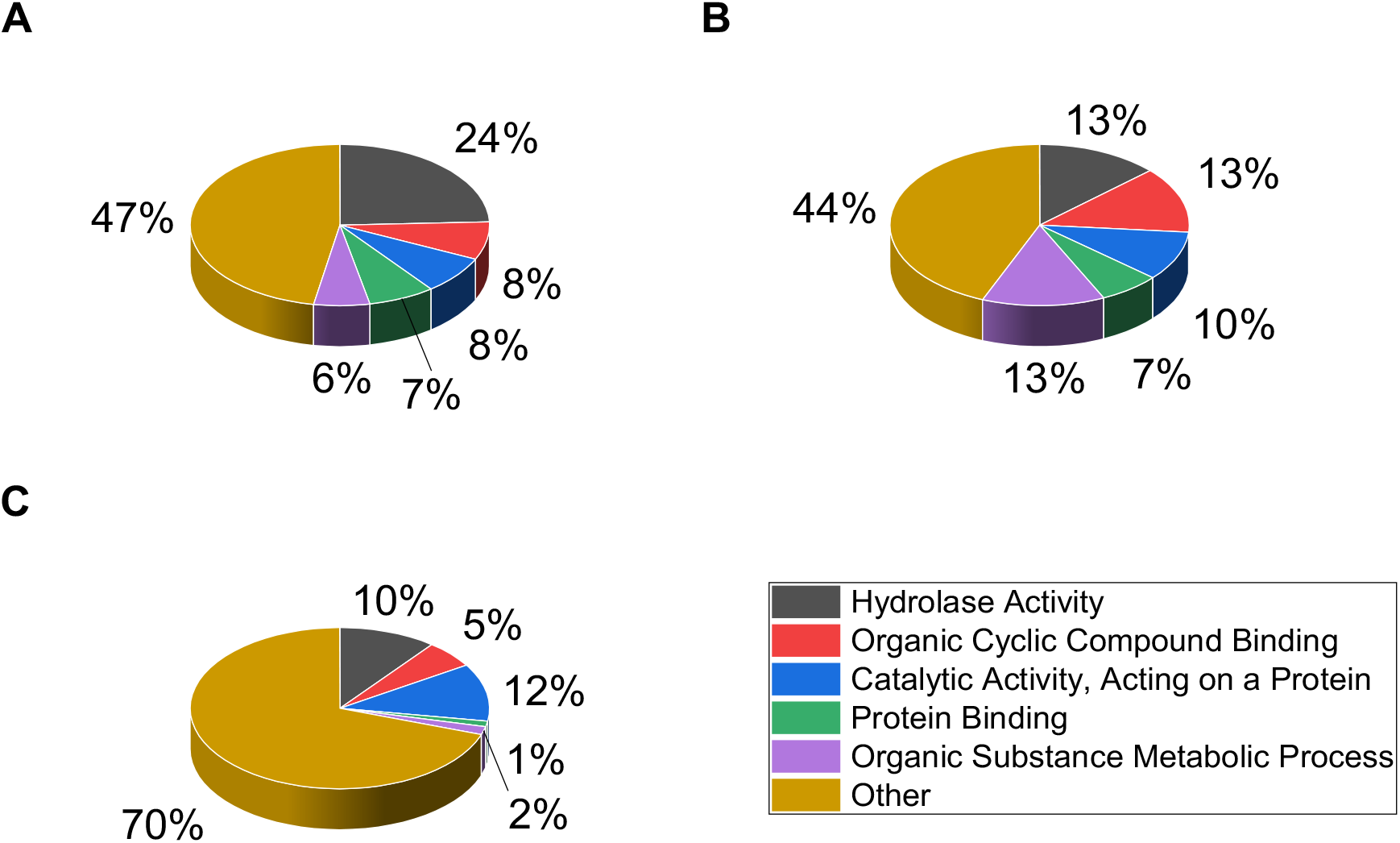
Function of protein complexes in the sets. The distributions of GO terms at depth 2 on the Gene Ontology tree are for proteins in (A) unbound decoy set, (B) model decoy set and (C) entire PDB.

## Conclusions

Scoring of protein-protein docking predictions generated by global docking scan is essential for docking methodologies. Development and validation of these methodologies are often performed on scring benchmark sets (docking decoys), which are supposed to be concise and nonredundant representations of the global docking scan output for a large and diverse set of protein-protein complexes. Two such protein-protein scoring benchmarks were built for the Dockground resource containing various datasets for the development and testing of protein docking methodologies. One set was based on the Dockground unbound docking benchmark 4, and the other set was based on protein models from the Dockground model-model benchmark 2. The decoys sets were designed to reflect the reality of the real-case scenario of docking applications (such as correct predictions as near-native rather than native structures), as well as to minimize applicability of trivial approaches not directly relevant to the development of scoring functions (reducing clustering of predictions in the binding funnel and disparity in structural quality of the near-native and non-native matches). The sets were further characterized by the source organism and the function of the protein-protein complexes. The sets represent a unique, user-friendly tool for the developing and testing of protein-protein scoring functions, and are freely available to the research community in the Dockground resource.

## Acknowledgments

This study was supported by NIH grant R01GM074255 and NSF grant DBI1917263.

## References

1. Vakser IA. Low-resolution structural modeling of protein interactome. Curr Opin Struct Biol. 2013;23:198–205.

2. Vakser IA. Protein-protein docking: From interaction to interactome. Biophys J. 2014;107:1785–1793.

3. Kundrotas PJ, Zhu Z, Janin J, Vakser IA. Templates are available to model nearly all complexes of structurally characterized proteins. Proc Natl Acad Sci USA. 2012;109:9438–9441.

4. Moal IH, Moretti R, Baker D, Fernandez-Recio J. Scoring functions for protein–protein interactions. Curr Opin Struct Biol. 2013;23:862–867.

5. Gromiha MM, Yugandhar K, Jemimah S. Protein-protein interactions: Scoring schemes and binding affinity. Curr Opin Struct Biol. 2017;44:31–38.

6. Lensink MF, Nadzirin N, Velankar S, Wodak SJ. Modeling protein-protein, protein-peptide, and protein-oligosaccharide complexes: CAPRI 7th edition. Proteins. 2020;88:916–938.

7. Vreven T, Moal IH, Vangone A, et al. Updates to the integrated protein-protein interaction benchmarks: Docking Benchmark Version 5 and Affinity Benchmark Version 2. J Mol Biol. 2015;427:3031–3041.

8. Kundrotas PJ, Anishchenko I, Dauzhenka T, et al. DOCKGROUND: A comprehensive data resource for modeling of protein complexes. Protein Sci. 2018;27:172–181.

9. Liu S, Gao Y, Vakser IA. DOCKGROUND protein-protein docking decoy set. Bioinformatics. 2008;24:2634–2635.

10. Gabb HA, Jackson RM, Sternberg MJE. Modelling protein docking using shape complementarity, electrostatics and biochemical information. J Mol Biol. 1997;272:106–120.

11. Lensink MF, Wodak SJ. Score_set: A CAPRI benchmark for scoring protein complexes. Proteins. 2014;82:3163–3169.

12. Chen R, Mintseris J, Janin J, Weng Z. A protein-protein docking benchmark. Proteins. 2003;52:88–91.

13. Mintseris J, Wiehe K, Pierce B, et al. Protein-protein docking benchmark 2.0: An update Proteins. 2005;60:214–216.

14. Hwang H, Pierce B, Mintseris J, Janin J, Weng Z. Protein–protein docking benchmark version 3.0. Proteins. 2008;73:705–709.

15. Hwang H, Vreven T, Janin J, Weng Z. Protein–protein docking benchmark version 4.0. Proteins. 2010;78:3111–3114.

16. Chen R, Li L, Weng Z. ZDOCK: An initial-stage protein-docking algorithm. Proteins. 2003;52:80–87.

17. Pierce B, Weng Z. ZRANK: Reranking protein docking predictions with an optimized energy function. Proteins. 2007;67:1078–1086.

18. Gray JJ, Moughon S, Wang C, et al. Protein–protein docking with simultaneous optimization of rigid-body displacement and side-chain conformations. J Mol Biol. 2003;331:281–299.

19. Anishchenko I, Kundrotas PJ, Tuzikov AV, Vakser IA. Protein models: The Grand Challenge of protein docking. Proteins. 2014;82:278–287.

20. Anishchenko I, Kundrotas PJ, Tuzikov AV, Vakser IA. Protein models docking benchmark 2. Proteins. 2015;83:891–897.

21. Singh A, Dauzhenka T, Kundrotas PJ, Sternberg MJE, Vakser IA. Application of docking methodologies to modeled proteins. Proteins. 2020;88:1180–1188.

22. Janin J. Assessing predictions of protein–protein interaction: The CAPRI experiment Protein Sci. 2005;14:278–283.

23. Ruvinsky AM, Vakser IA. Chasing funnels on protein-protein energy landscapes at different resolutions. Biophys J. 2008;95:2150–2159.

24. O’Toole N, Vakser IA. Large-scale characteristics of the energy landscape in protein-protein interactions. Proteins. 2008;71:144–152.

25. Katchalski-Katzir E, Shariv I, Eisenstein M, Friesem AA, Aflalo C, Vakser IA. Molecular surface recognition: Determination of geometric fit between proteins and their ligands by correlation techniques. Proc Natl Acad Sci USA. 1992;89:2195–2199.

26. Vakser IA. Protein docking for low-resolution structures. Protein Eng. 1995;8:371–377.

27. Anishchenko I, Kundrotas PJ, Vakser IA. Contact potential for structure prediction of proteins and protein complexes from Potts model. Biophys J. 2018;115:809–821.

28. Kundrotas PJ, Anishchenko I, Dauzhenka T, Vakser IA. Modeling CAPRI targets 110-120 by template-based and free docking using contact potential and combined scoring function. Proteins. 2018;86:302–310.

